# CRISPRs for strain tracking and its application to microbiota transplantation data analysis

**DOI:** 10.1101/344358

**Authors:** Tony J. Lam, Yuzhen Ye

## Abstract

CRISPR-Cas systems are adaptive immune systems naturally found in bacteria and archaea. Bacteria and archaea use these systems to defend against invaders, including phages, plasmids and other mobile genetic elements. Relying on integration of invader sequences (protospacers) into CRISPR loci (forming spacers flanked by repeats), CRISPR-Cas systems store genetic memory of past invasions. While CRISPR-Cas systems have evolved in response to invading mobile elements, invaders have also developed mechanisms to avoid detection. As a result of arms-race between CRISPR-Cas systems and their targets, the CRISPR arrays typically undergo rapid turnover of the spacers with removal of old spacers and acquisition of new ones. Additionally, different individuals rarely share spacers amongst their microbiome. In this paper, we developed a pipeline (called CRISPRtrack) for strain tracking based on CRISPR spacer content, and applied it to fecal transplantation microbiome data to study the retention of donor strains in recipients. Our results demonstrate the potential use of CRISPRs as a simple yet effective tool for donor strain tracking in fecal transplantation, and also as a general purpose tool for quantifying microbiome similarity.

## Introduction

The gut microbiome serves to provide a range of symbiotic functions, amongst which include metabolism, immune system development, and pathogen resistance [1]. While the gut microbiome plays an important role as a modulator of host health and disease, commensal colonizers are often succeptible to distruption which has been shown to be associated with the development of disease states [2, 3, 4]. One such example, is the persistant and recurrent *Clostridium difficile* infection (CDI) which is often induced by the treatment of antibiotics [5]. In an attempt to increase intestinal microbial diversity and re-establish a stable microbiome, fecal microbiota transplantation (FMT) has been applied to the treatment of recurrent CDI [6] and other gastrointestinal disorders [7]. The reported success rate of FMT based on thousands of patients with recurrent CDI is 90% following one or more FMT [8, 9]. Although restoration of a microbiota appears to have a positive effect against gut dysbiosis that is thought to exist in these patients, the exact mechanisms of FMT have yet to be fully elucidated [8]. In addition to gastrointestinal disorders, recent studies have shown promising applicaiton of FMT to treat other types of diseases, including Parkinson’s disease [10] and autism [11].

Bacteria and Archaea are continuously exposed to mobile genetic elements (MGE), such as phages and plasmids. While MGEs may provide adaptive advantages through horizontal gene transfer, they can also cause damage by disrupting a variety of the host’s regulatory functions [12]. To protect against invasive genomes which might cause harm to the host genome, prokaryotes utilize a variety of defence systems including the adaptive CRISPR-Cas immune systems [13]. CRISPR-Cas systems rely on integration of invader sequences (spacers) into CRISPR loci that act as a genetic memory of past invasions [13]. The CRISPR arrays comprise of spacer sequences which are flanked by repeats [14]. Processed CRISPR transcripts (crRNA) are then utilized as guides by Cas proteins to cleave complementary invader nucleic acids [15]. While prokaryotes have evolved CRISPR-Cas systems to target foriegn genetic elements, invaders have also evolved to evade CRISPR-Cas detection (i.e. localized protospacer adjacent motif [PAM] mutations) [16]. Although CRISPR arrays store information in regard to past encounters, spacer sequences of CRISPRs remain dynamic and often exhibit turnover between spacer sequences due to an arms race between CRISPR-Cas systems and its targets [14].

Since the discovery of CRISPR-Cas systems, much effort has been placed into understanding the mechanisms which govern the regulation of the spacers within CRISPR arrays. Erdmann and Garrett [17] demonstrated that three of the six CRISPR loci of *Sulfolobus solfataricus* rapidly acquire new spacer sequences from a conjugative plasmid present in a virus mixture, into the CRISPR arrays either adjacent to the leader sequence (which is typically found between the CRISPR array and the *cas* genes) or internally. LopezSanchez and collegues [18] studied the CRISPR-Cas systems in *Streptococcus agalactiae* and they found that a fraction of the *S. agalactiae* population with lost spacer sequences allowed for the transfer of a MGE in this subpopulation and a rapid response to altering selective pressures. Collectively, these studies reveal the dynamics which surround the maintenance of spacers within CRISPR arrays.

CRISPR-Cas systems have shown characteristics of spacer aquisition, as well as spacer turnover, which highlight the ever evolving nature of these systems. Similarly to bacteria found throughout the environment, microbial organisms found within the human microbiome (which mostly comprise of eubacteria) also carry CRISPR arrays which are dynamic in nature, with CRISPR locus expansion and contraction capabilities. With the ever changing composition of the CRISPR loci, gut microbiome found within human individuals often bear CRISPR arrays with unique spacer sequences. By taking advantage of the properties of spacer aquisition and retention within CRISPRs, the CRISPR spacers can potentially be used as molecular markers for typing and strain level species tracking purposes [19].

While the underlying dynamics and mechanisms of FMT remain largely undiscovered, efforts have been made to unvail these details by first understanding the impact of FMT at a microbial ecology level. To understand the effects of FMT induced microbiome reconstruction, it becomes important to understand the success of bacterial engraftment following fecal transplantation. Several studies have shown the success of utilizing single-nucleotide variants (SNV) based methods for tracking the dynamics between donor and recipient microbiomes following FMT[20, 21, 22]. The study by Li et al. on metabolic syndrome patients using SNV in metagenomes enabled the quantification of the extent of donor microbiota colonization after FMT, revealing extensive coexistence of donor and recipient strains, persisting 3 months after treatment. The authors also found that same-donor recipients displayed varying degrees of microbiota transfer, indicating individual patterns of microbiome resistance and donor-recipient compatibilities. Smillie et al. [22] developed StrainFinder, a tool to infer strain genotypes based on detected SNVs from FMT microbiomes and track strains over time. The successful usage of SNV based methods highlight the importance of understanding the microbioal ecology on a strain level. However, SNV calling in metagenomics can be complicated by the uneven abundance distribution of the bacterial species living in the same community and the coexistence of closely related species. Even worse, there is currently no strict definitions of what a bacterial or archaeal strain is [23].

Here we take advantage of the uniqueness of the CRISPR spacer contents in different human subjects and develop a method based on CRISPR spacer for the quantification of donor strain retention in FMT recipients. We utilize and leverage the tools we have developed for the identification and characterization of CRISPR-Cas systems in metagenomes. As compared to SNVs, spacers are relatively large entities of approximately 20 to 50 bps long, so they are more straightforward to characterize. Applying our tool, CRISPRtrack, to two fecal microbiota transplantation datasets demonstrate the potential use of CRISPRs as a simple yet effective tool for donor strain tracking in fecal transplantation.

## Methods

### Identification of CRISPR arrays in genomes and metagenomes

In this study, we utilize two approaches we have previously developed for the identification of CRISPR arrays from genomes and metagenome assemblies: a reference-based and *de novo* based approach. While our previous study [24] has shown that in some instances CRISPR arrays are difficult to assemble from shotgun metagenomic sequencing data due to the presence of repetitive regions in CRISPR arrays and may lead to under-representation of CRISPRs. We found that metagenome specific assemblers such as MEGAHIT [25] have since provided improved assemblies for identification of CRISPR-Cas systems comparatively to previous assemblers [26]. As a result, in this study we utilize MEGAHIT to assemble metagenomes using k-mer size parameter (k-list = 21,41,61,81,99). Contig assemblies were then used for downstream CRISPR-Cas identification.

For *de novo* based prediction, we utilize CRISPRone a tool we previously developed [27] for the prediction of CRISPR-Cas systems. CRISPRone utilizes metaCRT, a modified variant of CRT, for the identification of novel CRISPR arrays [24]. The use of metaCRT exploits the structure of CRISPR arrays and searches for sequences containing repeat-spacer like structures. CRISPRone applies additional filters to remove suspicious CRISPR arrays through various methods such as the identification and removal of tandem repeats and STAR-like elements [28, 29].

In our reference based approach, we use a set of CRISPR repeat sequences associated with human gut microbiomes to be used as reference repeats. Utilizing this set of reference repeats with CRISPRAlign [24], we are able to identify CRISPRs which share similar repeats as our reference repeats. Comparatively to *de novo* approaches, reference based methods hold an advantage by using a set of carefully identified reference repeats which reduces the chances of false positive CRISPR arrays. Contrastingly, as reference based approaches only searches for CRISPRs sharing similar reference repeats, reference based methods may result in a underrepresented set of CRISPRs by missing any CRISPRs which may have dissimilar CRISPR repeats.

### Quantification of the existence of donor-species in recipient using CRISPR spacer contents

Using predicted CRISPR arrays, all corresponding putative CRISPR spacers are extracted. Utilizing the set of all extracted spacers, cd-hit-est (-c 0.9) is used to cluster spacers into their representative groups. Each cluster represents a unique spacer, and all spacers grouped into the same cluster are considered *the same,* to allow mismatches in the spacers and potential sequencing errors. Donor species presence within recipient samples at a given timepoint (rt), can be quantified by computing the sharing of spacers between the recipient microbiome and donor microbiome as following:
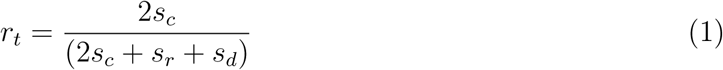
where *s_c_* denotes the number of clusters containing spacers from both the recipient and the donor sample, *s_r_* denotes the number of clusters containing only spacers from the recipient, and *s_d_* denotes the number of clusters containing only spacers from the donor.

While CRISPR-Cas containing isolates may provide a unique set of spacers, isolates from different individuals can sometimes share common spacers. To normalize for spacers shared between donor and recipient prior to fecal transplantation (day 0), the common subset of shared spacers between donor and recipient samples may be removed from the downstream analysis.

### The fecal transplantation datasets and the HMP datasets

We utilized three sets of previously published datasets to test our new method. For clarity, we denote these datasets as FMT-Li [21], FMT-Smillie [22], and HMP [30]. FMT-Li datasets [21] were downloaded from the European Nucleotide Archive (ENA) under the accession number PRJEB12357. The FMT-Li datasets include metagenomic sequencing data from five patients (FMT1-FMT5) receiving microbiota transplantation from three healthy donors (Don1-Don3), in which stool samples were collected from patients at multiple time points spanning between pre-treatment to 84 days post-treatment. FMT-Smillie datasets [22] were downloaded from the ENA under the accession number PRJEB23524. The FMT-Smillie datasets consist of microbiome sequencing data obtained from four donors, and 19 patients whose stool samples were collected at multiple timepoints spanning between pre-treatment to 135 days posttreatment. HMP datasets were published in [30], and were derived from the dacc website at https://dacc.org. We used a total of 95 stool microbiome datasets from the HMP collection.

### Availability of the software

We implemented a package called CRISPRtrack for identification of CRISPRs in metagenome assemblies, and quantification of the retention of donor species in recipients. The package is available for download at sourceforge at: https://sourceforge.net/projects/crisprtrack/. The package contains a few tools that we previously developed and additional scripts that we developed for this study for computing the sharing of the CRISPR spacers. The package also includes tools for further analyses and visualizations of the results. CRISPRtrack supports both approaches for characterizing CRISPRs: the reference based approach using gut microbiome-related reference repeats (called CRISPRtrack-ref) and *de novo* prediction by CRISPRone (CRISPRtrack-denovo). The package outputs spacer-subject tables, similarity scores between microbiomes based on spacer content sharing, and tracking plots of donor strains in recipients based on CRISPR spacer sharing.

## Results

We first report the identification of reference CRISPR repeats from human gut-associated bacterial genomes. We then study the baseline similarity of the spacer contents between human individuals. Finally we report the results of applying CRISPRtrack to two FMT datasets (FMT-Li and FMT-Smillie).

### CRISPRs in common gut microbial species

We identified CRISPR-Cas systems in common gut microbial species, with the goal of defining a set of reference CRISPR repeats for reference based identification of CRISPR spacers in gut microbiomes. We first checked 42 common strains found in the fecal microbiota samples [21]. Also in our previous study [26], we analyzed a different cohort of gut microbiome datasets [31], from which we were able to identify 33 unique CRISPR repeats. Combining the two subsets of CRISPR repeats, we were able to compile a collection of 64 unique CRISPR repeats to be used as reference repeats. The sequences of the reference repeats are included in the CRISPRtrack package.

We show in Figure 1, a subset of examples of CRISPR-Cas systems found in these common gut-associated bacterial species. Unsurprisingly, most of the CRISPR-Cas systems found in the gut-associated microbial genomes belong to the type I CRISPR-Cas system. Additionally, we found type II, III, and V CRISPR-Cas systems in these genomes.

**Figure 1:**
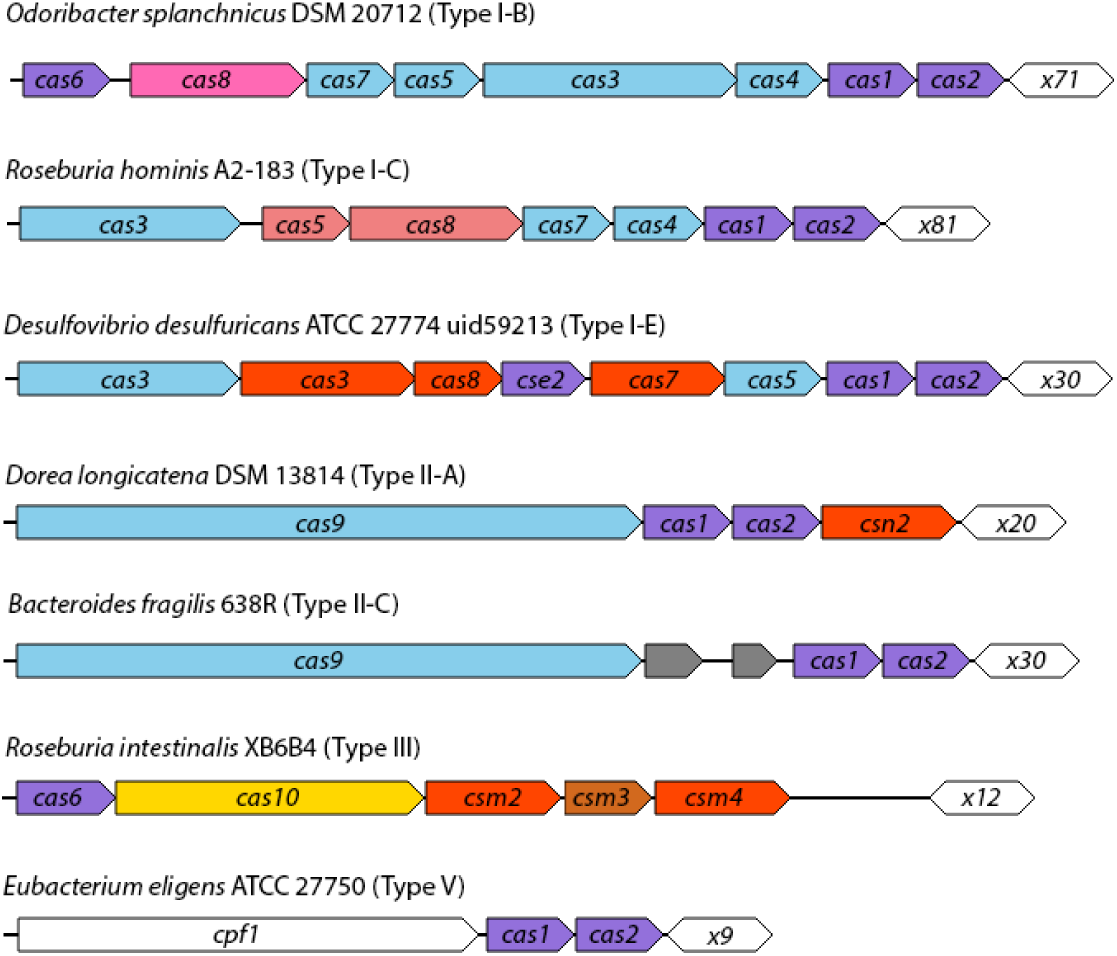
Representative CRISPR-Cas systems found in common human gut-associated bacterial species. The arrows in different colors represent the *cas* genes, and the open hexagons represent the CRISPR repeat-spacer arrays, with numbers following the letter *x* indicating the number of spacers found in each array.

### Spacer sharing between human individuals (the baseline)

The HMP dataset involves 95 microbiomes derived from 79 human subjects, with some subjects each having two microbiomes samples. We applied both reference based and *de novo* approaches to identify CRISPR spacers in the HMP datasets, and used the spacers to estimate the baseline of the spacer sharing between different individuals.

The reference based approach resulted in a total of 26,074 spacer clusters, which were then used to compute the spacer content similarity. We show that gut microbiomes from different individuals share significantly fewer spacers (with the median spacer content similarity = 0.0049) as compared to gut microbiomes from the same individual but different time points (with the median spacer content similarity = 0.66) (Figure 2A). To further study spacer sharing among unrelated individuals, we focused our analysis on individuals who had greater than one sample. This reduced the number of spacer clusters to 23,868, among which, 19,634 clusters (82%) contained only a single spacer indicating the spacer is unique to the individual. From the spacer clusters, only 63 clusters (0.26%) were found to be shared among 10 or more individuals. Through utilizing this analysis, we were able to show that while some spacers are shared among individual samples, a large portion of spacers remain unique to individual samples and are not shared with others. Spacers shared by many individuals are likely to originate from inactive CRISPR arrays, which do not exhibit active turnover of spacers.

Similarly, we analyzed the CRISPR spacer sharing between individuals using spacers predicted through the *de novo* approach. The *de novo* approach predicted many more CRISPR spacers as compared to the reference based approach, resulting in a total of 48,204 spacer clusters. The difference is expected as the reference based approach only identified CRISPR arrays containing repeats similar to the reference repeats, whereas the *de novo* approach identified *all* CRISPR arrays. Reassuringly, analysis of *de novo* predicted spacers revealed consistent results that microbiomes from different individuals shared few CRISPR spacers whereas the microbiomes of the same individual shared substantially more spacers, as seen in Figure 2B.

**Figure 2:**
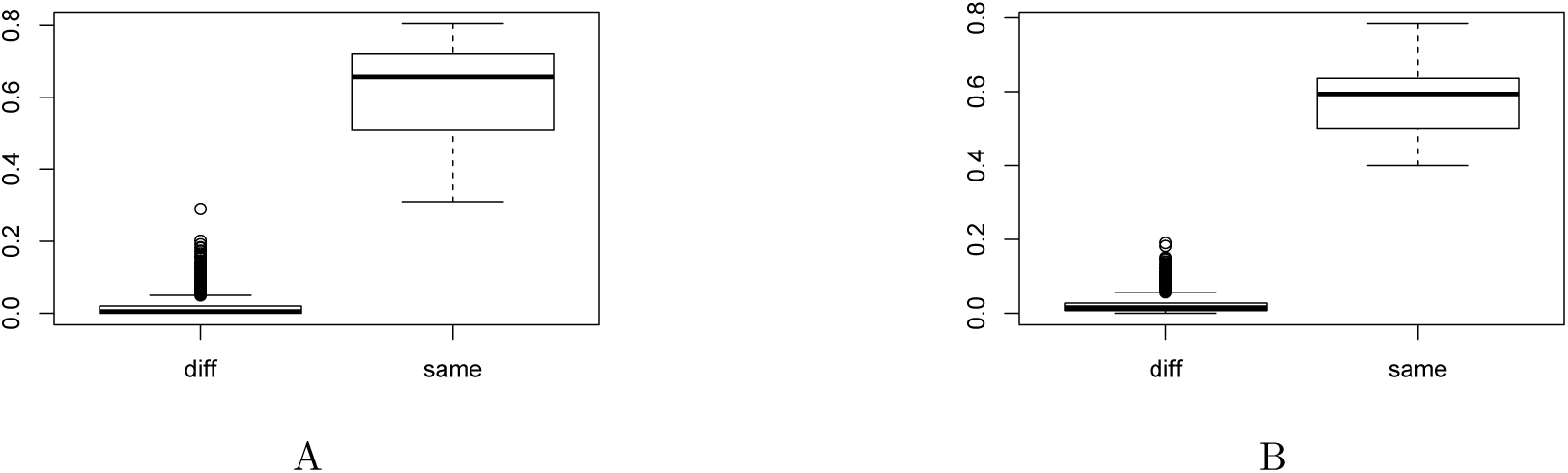
Sharing of the CRISPR spacers among HMP individuals. These two figures show the boxplots of spacer content similarity between microbiomes from different individuals (diff) and microbiomes from the same human subject (same). (A) is based on spacers identified using the reference based approach, and (B) is based on *de novo* identification of CRISPR arrays.

By obtaining the distribution of spacer similarity between different HMP individuals, we were able to calculate the 95th percentiles for CRISPR spacer content similarities. The 95th percentiles of CRISPR spacer content similarity between different HMP individuals were 0.062 and 0.056, for spacers derived by the reference based approach and *de novo* approach, respectively. These 95th percentiles were then used as the baseline CRISPR spacer similarities for checking the sharing of spacers between FMT recipients and their donors as shown in Figure 3 and Figure 4 below.

**Figure 3:**
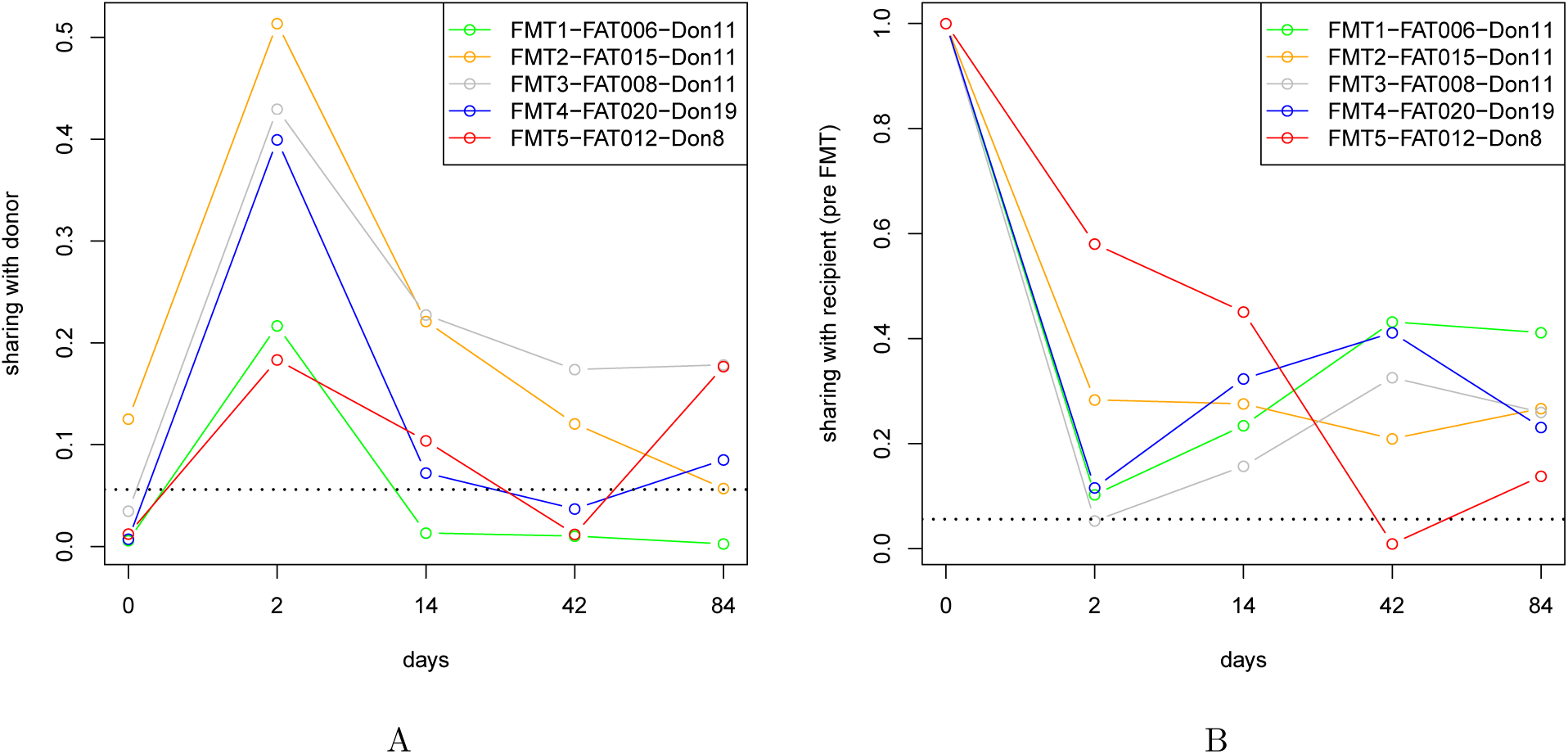
Tracking of donor spacers and recipient’s own spacers over time after FMT. (A) shows the CRISPR spacer similarity between the recipients and their corresponding donors. (B) shows the CRISPR spacer similarity between the recipients after FMT and their pre-FMT counterparts. Lines connect time point samples from the same individual. The time axis (i.e., the x-axis) was not scaled for clarity. The dotted black lines in the plots indicated the 95th percentiles of the spacer similarities between different individuals, inferred from the HMP datasets.

**Figure 4:**
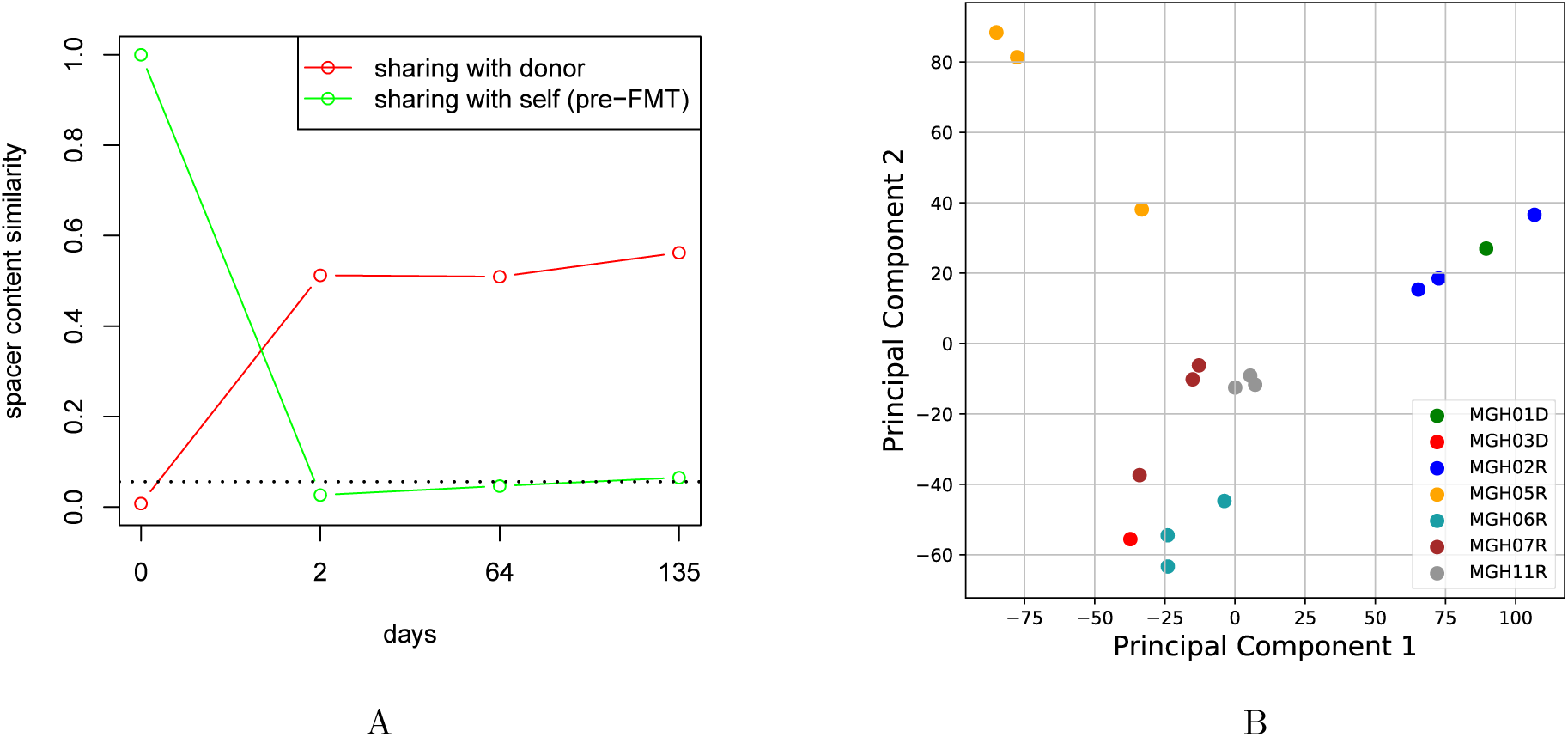
CRISPRtrack results for the FMT-Smillie dataset. (A) Tracking of the donor-and pre-FMT recipient-specific spacers in recipient MCH02R. (B) shows the clustering of microbiomes according to their CRISPR spacer sharing, after applying the PCA analysis.

### Application of CRISPRtrack to FMT-Li dataset

We applied CRISPRtrack to characterize the CRISPR spacers from the donor and recipient microbiome data (FMT-Li) [21], and quantified the retention of donor CRISPR spacers in recipients using the CRISPR spacer content. The spacer similarity plots (Figure 3; based on spacers predicted *de novo)* show that the recipient microbiome contains similar CRISPR spacers as the donor microbiome, especially during the early timepoints post-FMT, indicating that a significant amount of donor-sourced bacteria were transferred into the recipient and retained for that period of time. Our results also show that after significant reduction of the recipient’s own bacteria (as indicated as the low spacer similarity between recipient and its pre-FMT microbiome), there is a resurgence of the recipient’s original strains (Figure 3B). We note using spacers predicted by the reference-based approach showed consistent results with the *de novo* based approach.

Below we highlight a few comparison of our results with the results from the previously reported SNV-based analysis [21].

- It was mentioned that marked differences in colonization success were observed between allogenic recipients who shared a donor (FMT1, 2, and 3). Three months after treatment, FMT2 and 3 retained a higher amount of donor-specific SNVs compared with FMT1 (46.1%, 56.6%, and 12.0%, respectively). CRISPRtrack revealed a similar trend: FMT1 retained the fewest donor-specific spacers, whereas FMT2 and FMT3 retained more.
- The SNV based analyses showed the highest retention of donor strains in FMT 2 and FMT 3; CRISPRtrack revealed the same trend.
- The SNV based analyses showed the resurgence of donor-specific strains in FMT5 after day 14; our analysis showed similar trend, although the resurgence peaked on day 84 instead of day 42.

In our analysis, we observe that FMT2 (prior to treatment, i.e., day 0) shares relatively more common CRISPR spacers with the donor microbiome comparatively to other recipients (see Figure 3A). We also examine the spacers identified using the reference-based approach to study the spacers that are common to donor and recipient. For example, CRISPR arrays containing repeats that are almost identical (with one bp difference) to the repeat identified in the *Bacteroides fragilis 638R* genome (named BfragL47-II), are found to share five spacers. The sequence of BfragL47-II is GTTGTGATTTGCTTTCAAATTAGTATCTTTGAACCATTGGAAACAGC, and at position 31 the repeat identified in the donor and recipient CRISPRs have T instead of A. Another example is CRISPRs arrays containing repeats that were identical to the CRISPR repeat found in reference genome *Alistipes shahii* WAL8301 (named AshahL36 whose sequence is GTTGTGGTTTGATGTAGAATTTCGATAAGATACAAC). Interestingly, the CRISPR identified in *A. shahii* is an orphan CRISPR without *cas* genes. An array containing AshahL36 repeats was assembled from the recipient dataset, which contains 9 spacers, a subset of the 36 spacers in a much longer array assembled from the donor dataset.

### Application of CRISPRtrack to FMT-Smillie dataset

We applied CRISPRtrack to a more recent FMT microbiome dataset (FMT-Smillie) [22]. Although this dataset involved more human subjects than the earlier mentioned FMT dataset (FMT-Li) [21], many recipients only contained one microbiome sample. Here we focus on the individuals that have multiple microbiome samples post-FMT. We also note that the samples were collected at different time intervals, unlike the FMT-Li dataset which sampled all recipients at regular time intervals. Another major difference between FMT-Smillie and FMT-Li datasets is that most recipients in the FMT-Smillie study received antibiotic treatments prior to FMT.

As an example, Figure 4A shows the tracking of *de novo* predicted CRISPR spacers in samples from recipient MGH02R. The patient’s pre-FMT microbiome apparently contained fewer bacterial species as compared with post-FMT microbiomes: the pre-FMT microbiome assembly contained approximately 85 Mbp, whereas post-FMT (day 2) microbiome assembly contained approximately 396 Mbp. As a result, isolates from the FMT-Smillie dataset exhibit few commonly shared spacers between pre-FMT and post-FMT samples (Figure 4A). By contrast, the pre-FMT and post-FMT microbiomes in the FMT-Li dataset share significantly more spacers, which was likely due to FMT-Li patients not receiving antibiotic treatment prior to FMT (Figure 3).

Focusing on the data involving two donors (MGH01D, and MGH03D) from the FMT-Smillie dataset, we performed a two dimensional principal component analysis (PCA) on the spacer-subject table produced by CRISPRtrack. Unsurprisingly, the PCA revealed clusters of samples involving the same recipient and the same FMT donor (see Figure 4B). We note for the same reason (antibiotic treatment prior to FMT) that pre-FMT microbiomes have very low bacterial diversity, we did not include the pre-FMT samples in this analysis. In Figure 4B, donor samples are depicted in green (MGH01D) and red (MGH03D). Recipient samples MGH06R (azure blue), MGH07R (brown), and MGH11R (grey) received FMT from MGH03D donor samples and are shown to cluster together. Similarly, MGH02R (blue) which received samples from donor MGH01D (green) are shown to cluster together. However, MGH05R (orange) which received FMT from MGH03D shared the least CRISPR spacers comparatively to its donor.

## Discussion

Our study shows the potential of using CRISPRs for tracking the engraftment of CRISPR containing donor strains in recipients following FMT. CRISPRtrack provides two approaches for spacer identification: reference and *de novo* based. We expect that the reference-based approach will be suitable for studying microbiota from well-studied environments (i.e. gut microbiome), as this approach relies on the curation of reference CRISPR repeats. Alternatively, the *de novo* approach will be simpler to apply for less well-studied microbiota data analysis. In terms of running time, the reference based approach is slower than the *de novo* approach, since the reference based approach involves an alignment step to find segments in metagenomic assemblies that are similar to reference CRISPR repeats. Still, both approaches are fast: using only a single process (Intel Xeon CPU E5-2623 v3 @ 3.00GHz), the whole pipeline completed in 27 and 217 minutes for the *de novo* and reference based approaches, respectively, on the FMT-Li collection, and the time was 32 and 342 minutes for the *de novo* and reference based approaches, respectively on the FMT-Smile collection. We note that although various methods, including the method reported here, have been developed for the tracking donor strains in FMT recipients, predictive models used for the prediction of successful FMT based on microbiome data still remain lacking [21, 22].

CRISPRs provide a unique advantage in that they can provide a unique subset of spacers which can be utilized as molecular markers, providing a high resolution approach to differentiate bacterial strains from separate individuals. While we show that CRISPR based tracking methods hold the potential of revealing microbial community dynamics, we also acknowledge the limitations of such approaches. The fast evolving nature of CRISPRs causes constant spacer acquisition and turnover which limits its use to short term tracking. Additionally, not all prokaryotes contain CRISPR-Cas systems, thus CRISPR spacers cannot be used for the tracking of microbial strains which lack CRISPR-Cas systems. With these potential limitations of using CRISPR spacers in mind, we believe that CRISPR spacers can still be used to serve as sensitive molecular markers for tracking microbes, especially when we consider microbial communities as a whole.

Utilizing the methods we have developed for using CRISPRs to track donor strain retention, we can begin to consider exploring further questions about the dynamics of CRISPR spacers in FMT patients. Potential avenues of exploration may include the dynamics of CRISPR spacer turnover following FMT, as well as understanding if spacer acquisition in recipient CRISPRs can be correlated to donor microbiomes post-FMT.

